# Velociraptor: Cross-Platform Quantitative Search Using Hallmark Cell Features

**DOI:** 10.1101/2024.05.01.591375

**Authors:** Claire E. Cross, Cass Mayeda, Stephanie Medina, Madeline J. Hayes, Saara Kaviany, James A. Connelly, Jeffrey C. Rathmell, Kyle D. Weaver, Reid C. Thompson, Lola B. Chambless, Rebecca A. Ihrie, Jonathan M. Irish

## Abstract

A key challenge for single cell discovery analysis is to identify new cell types, describe them quantitatively, and seek these novel cells in new studies often using a different platform. Over the last decade, tools were developed to address identification and quantitative description of cells in human tissues and tumors. However, automated validation of populations at the single cell level has struggled due to the cytometry field’s reliance on hierarchical, ordered use of features and on platform-specific rules for data processing and analysis. Here we present Velociraptor, a workflow that implements Marker Enrichment Modeling in three cross-platform modules: 1) identification of cells specific to disease states, 2) description of hallmark features for each cell and population, and 3) searching for cells matching one or more hallmark feature sets in a new dataset. A key advance is that Velociraptor registers cells between datasets, including between flow cytometry and quantitative imaging using different, overlapping feature sets. Four datasets were used to challenge Velociraptor and reveal new biological insights. Working at the individual sample level, Velociraptor tracked the abundance of clinically significant glioblastoma brain tumor cell subsets and characterized the cells that predominate in recurrent tumors as a close match for rare, negative prognostic cells originally observed in matched pre-treatment tumors. In patients with inborn errors of immunity, Velociraptor identified genotype-specific cells associated with *GATA2* haploinsufficiency. Finally, in cross-platform analysis of immune cells in multiplex imaging of breast cancer, Velociraptor sought and correctly identified memory T cell subsets in tumors. Different phenotypic descriptions generated by algorithms or humans were shown to be effective as search inputs, indicating that cell identity need not be described in terms of per-feature cutoffs or strict hierarchical analyses. Velociraptor thus identifies cells based on hallmark feature sets, such as protein expression signatures, and works effectively with data from multiple sources, including suspension flow cytometry, imaging, and search text based on known or theoretical cell features.

## Introduction

A cell can be classified based on its morphology, location, protein expression, or a combination of these features ^1^. The method of cell identification depends on the technology used to measure cellular features. In the field of flow cytometry, the classic approach to identify cells is manual biaxial gating, which uses strict thresholds of marker positivity and negativity that are used to distinguish cell populations ^2,3^. Hierarchical filtering for populations using manual gating excludes cells that don’t perfectly match a phenotype of interest, even if a cell falls just short of a threshold for a single protein. This approach is subjective and labor intensive ^4^, especially when analyzing high-parameter mass cytometry datasets that can contain measurements of greater than 40 markers ^5,6^. Additionally, manual gating is biased towards identification of well-established populations that have known patterns of protein expression ^7^.

Unsupervised machine learning cell identification approaches aim to overcome the limitations of subjectivity and bias towards known populations that are endemic to manual biaxial gating. Cell populations can be distinguished in low-dimensional projections of high-dimensional data, which includes all cells and not just known populations ^4,7,8^. To identify different populations within a sample, unsupervised clustering methods, such as k-means clustering and FlowSOM ^9^, group cells into a pre- defined number of non-overlapping subpopulations that ideally represent distinct cell types. However, these disjoint clustering methods have often been limited by the requirement of *a priori* knowledge of the number of distinct populations that exist in a sample. Additionally, stochasticity can result in varying assignment of cell types across separate analyses. The resulting disjoint clusters can also contain heterogeneous mixtures of cells that belong to distinct cellular states that should not be grouped into a single phenotype. Alternatively, we and others have established the use of cell-specific approaches that utilize overlapping local phenotypic neighborhoods to identify populations ^10–12^. This local phenotypic neighborhood approach continuously tracks phenotypic space, which enables detection of rare cells (<5% of a sample) and subtle phenotypic shifts that could have been overlooked if that cell were to be placed into a larger, disjoint cluster ^10^. Individual populations can then be tested for association with external variables, such as infection status or overall survival time, using various unsupervised machine learning algorithms ^13–16^.

Historically, identifying the phenotype of a subpopulation of cells that is associated with an external variable has required manual review and expert annotation of markers that are expressed by that group of cells. The development of Marker Enrichment Modeling (MEM) enabled automatic quantification of population phenotypes ^17^. MEM generates a label that includes the markers that are enriched in different populations as well as the relative level of enrichment of each marker. These labels are quantitative and both human- and machine-readable, meaning that not only is a population’s identity summarized in a way that can be interpreted by a scientist, but a MEM label can also be used as the starting point for subsequent computational analyses. For example, the phenotypes of multiple populations can be compared by computing a ΔMEM label that quantifies pairwise differences in feature enrichment between two populations ^18^. Thus, cell identity can be quantified based on the expression or enrichment of various features, such as proteins. However, a historical lack of standardization and use of differing approaches to define cell identity across fields has led to the generation of multiple independent methods to identify the same population of cells ^1^. Even in a well-established field such as immunology, different groups may use different features to identify the same cell population (e.g., using CD127, CD25, FOXP3, or a combination of these features to identify regulatory T cells). Thus, identification of similar populations in additional datasets based on a learned phenotype still requires a high degree of manual supervision to identify cells that may have been characterized using different features in order to validate findings.

Populations of interest identified in analyses should be validated for stability and generalizability using approaches such as k-fold cross validation and leave-one-out cross validation ^19,20^. A stable cluster will be composed of the same, or similar cells, across multiple runs of a machine learning pipeline on repeated samples of the dataset. Population identification in newly collected samples will reveal the generalizability of initial results. The gold standard to biologically validate a result is to use a different platform to ensure the observed population is not a technological artefact. However, there are many challenges in validating automatically identified cell populations across experiments and modalities. Inherent technological differences cause single cell platforms to collect data that can span widely different ranges. As a result, a common cross-platform cell identification approach is to develop a manual gating scheme to select cells based on marker positivity and negativity ^10,16^. Computational algorithms have also been developed to integrate distinct data types by combining the data into a common phenotypic space ^21–23^. However, these methods require (1) a large overlap between marker panels as they were developed to integrate scRNAseq data, (2) extensive data pre-processing steps correct for technological differences and to transform data onto similar scales, and often (3) a “matchability test” to ensure that the two datasets are amenable to a joint analysis ^24^. While a single pooled analysis better captures the natural variability that exists among patients, the number of cells analyzed per patient is often limited to the fewest number of cells collected from a donor to prevent a single sample from dominating an analysis ^14,25–27^. This equal sampling limits the total number of cells analyzed, which can reduce the overall power of the analysis. Another limitation of a pooled analysis can be that upon collection of an additional sample to be included in the cohort, samples must be re- analyzed. Algorithms such as UMAP and SCAFFOLD ^8,28^ allow additional samples to be added into a previously generated low dimensional embedding but assume that there are no sample-specific cell types or patterns of expression that were not present in the original analysis. Ideally, a cross-dataset identification algorithm will require minimal overlap between feature panels, analyze samples individually rather than performing a pooled analysis in a common phenotypic space, and control for potential differences in data scales due to minor batch effects or different data types.

Here, we present Velociraptor, a novel cell identification workflow that reveals clinically relevant cell populations, identifies the phenotypes of those populations, and seeks out similar cells across experiments and single cell platforms. We use Velociraptor to address biological challenges that are regularly encountered across diverse settings of disease, including automated cell identification in cases of limited sample or cell numbers and matching of cell populations across technologies. The discovery and seeking of populations across datasets ultimately allowed for identification of rare and abundant cells associated with external variables, determination of the most essential proteins that distinguish a population from other cells, and automated validation of cell subsets across single cell platforms without forcing distinct data into a common phenotypic space.

## Results

### Velociraptor overview

The Velociraptor workflow consists of two novel cell identification tools that can be used in tandem or individually: Velociraptor-Claw (VR-Claw) and Velociraptor-Eye (VR-Eye) (**Figure 1A**). VR-Claw utilizes local phenotypic neighborhood identification to reveal cells that are associated with continuous variables (e.g., overall survival time). MEM quantifies the phenotype of a population of interest, and that MEM label is used to seek similar cells in new samples with VR-Eye. This cell identification workflow addresses four biological challenges: 1) de novo cell identification and alignment with external labels, 2) cell identification and patient classification in new samples, 3) identifying cells across batches collected using the same platform, and 4) identifying cells across single cell platforms (**Figure 1B**). Each analysis was statistically validated using methods such as k-fold cross validation, whole sample analysis, repeated down-sampling, and leave-one-out cross validation (LOOCV) (**Figure 1C**). VR identified clinically relevant cell populations across experiments and single cell platforms (**Figure 1D**) as described in the analyses below.

**Figure 1.**
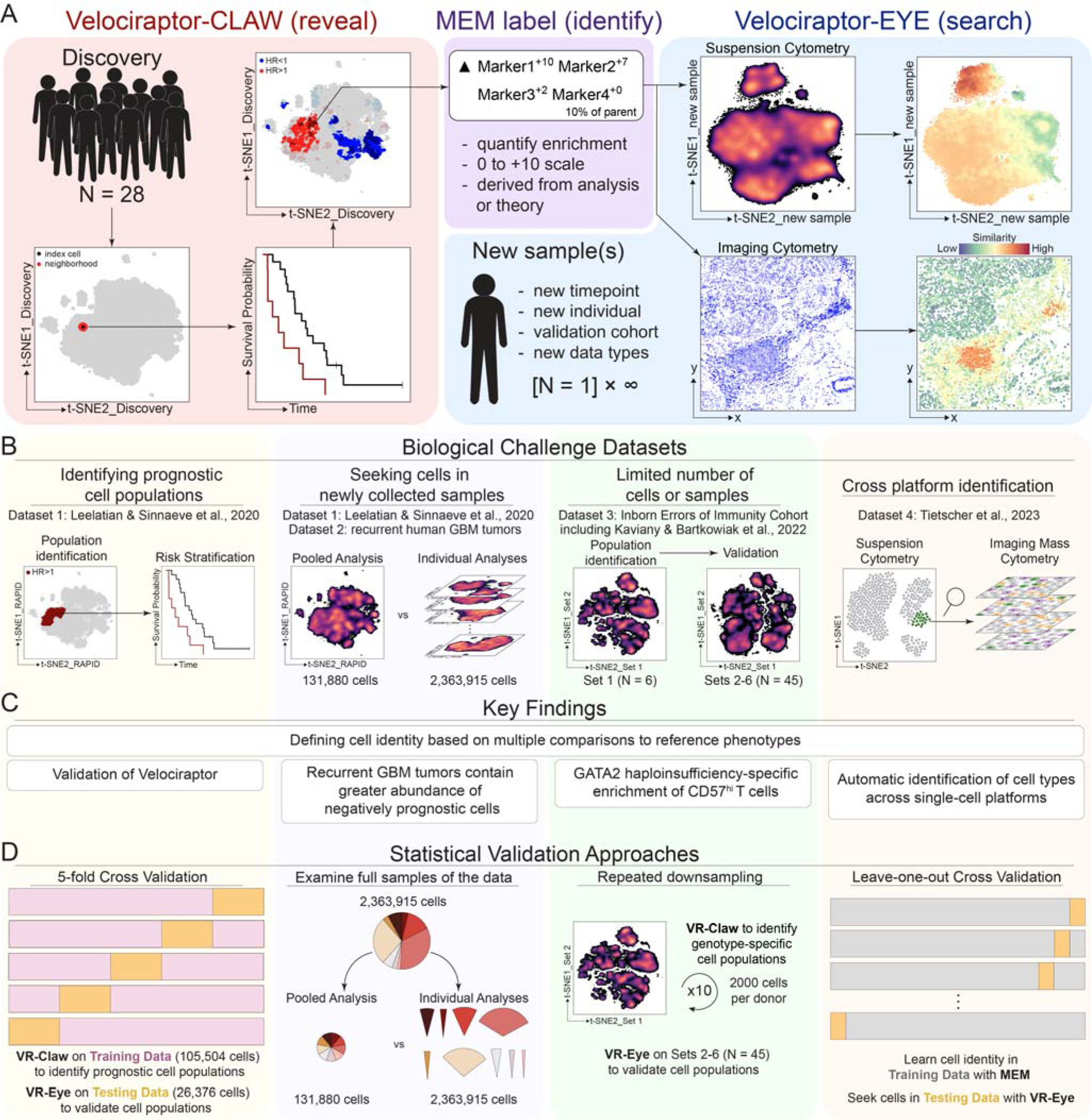
Overview of Velociraptor. **A)** The Velociraptor workflow includes two novel cell identification tools: (1) Velociraptor-Claw (VR-Claw), which reveals condition-specific cell populations (e.g., cells associated with overall survival), and (2) Velociraptor-Eye (VR-Eye), which seeks cells based on a user-defined phenotype. VR-Claw utilizes local phenotypic cell neighborhoods to identify clinically relevant cell populations. The upper right panel in the red box highlights cells associated with shorter (red) or longer (blue) survival times. Marker Enrichment Modeling (MEM) ^17^ can be used to automatically quantify population phenotypes, and MEM labels can then be used as input to VR-Eye. VR-Eye quantifies and plots similarity to a specified phenotype of interest (i.e., with a MEM label) with purple indicating low similarity and red indicating high similarity. **B)** Biological challenges and datasets explored in this manuscript. **C)** Key findings from Velociraptor for each biological challenge. **D)** Statistical validation approaches used to validate findings include k-fold cross validation, interrogation of cells not used to learn a phenotype of interest, repeated down-sampling, and leave-one-out cross validation.

### Velociraptor identified clinically relevant cell populations in primary GBM tumors

#### Velociraptor-Claw revealed prognostic GBM populations

The goal of the first analysis was to validate that Velociraptor could accurately identify cell populations that were known to be associated with overall survival times. The algorithm was tested using a published 36-dimensional mass cytometry dataset that characterized a cohort of 28 primary human glioblastoma tumors (**<u>Dataset 1</u>**) ^14^. This dataset has been shown to contain two distinct prognostic cell populations: 1) Glioblastoma Negative Prognostic (GNP) cells that co-express astrocytic marker S100B and stem-like marker SOX2 and are associated with shorter overall survival times (median survival of 144 days), and 2) Glioblastoma Positive Prognostic (GPP) cells that are characterized by high expression of EGFR and are associated with longer overall survival times (median survival of 836 days).

VR-Claw identified one stable GNP population (p<0.05, HR>1) and four stable GPP populations (p<0.05, HR<1; **Figure 2A**). Here, stable means that a similar cluster was significantly (p<0.05) associated with overall survival in each of the 5 rounds of cross validation. GNP cells in cluster 1 co-expressed S100B and SOX2 and displayed basal phosphorylation of multiple signaling proteins (MEM: S100B^+6^ SOX2^+5^ p-STAT3^+4^, CyclinB1^+3^ p-STAT5^+3^, p-S6^+3^, p-AKT^+2^, and p-NFkB^+2^; **Figure 2B**). GPP cells in clusters 2, 3, 4, and 5 occupied four distinct expression profiles (**Figure 2B**). Cluster 2 expressed a high level of EGFR (MEM: EGFR^+7^ SOX2^+3^ GFAP^+3^ p-NFkB^+2^ CD44^+2^). Cluster 3 co-expressed astrocytic markers S100B and GFAP as well as EGFR (MEM: S100B^+3^ EGFR^+3^ GFAP^+2^). Cluster 4 exhibited high basal signaling and lacked expression of canonical neural cell surface markers (MEM: p-AKT^+10^ p-STAT1^+3^). Cluster 5 displayed high levels of phosphorylation of p- STAT5 and p-NFkB along with EGFR expression (MEM: p-STAT5^+9^ EGFR^+5^ p-NFkB^+4^ S100B^+2^ CD56^+2^ SOX2^+2^ CD44^+2^ GFAP^+2^).

**Figure 2.**
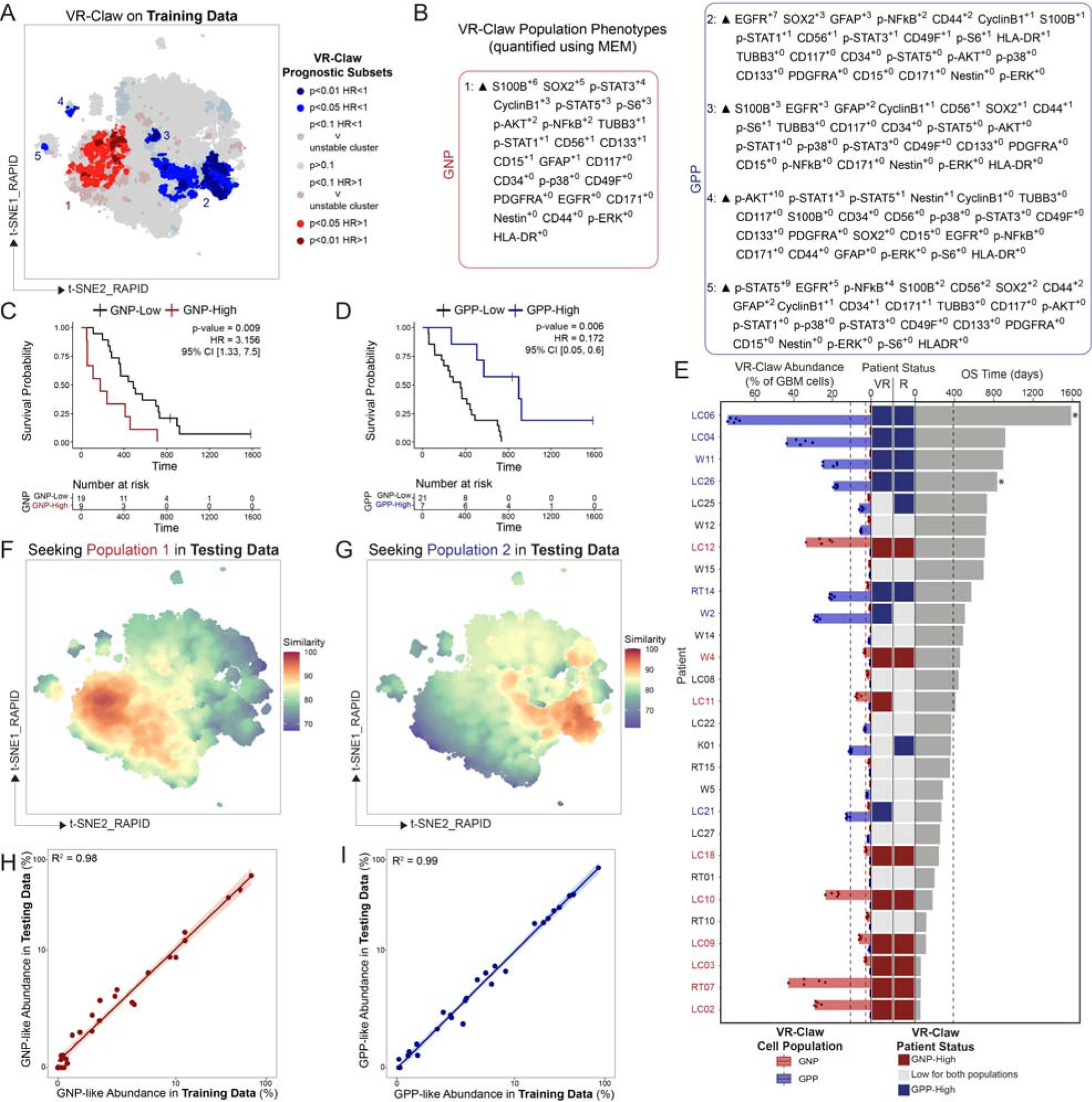
Velociraptor refines GNP and GPP cell identification and patient classification. **A)** VR-Claw analysis of CD31^-^CD45^-^ cells from the RAPID cohort. Cell neighborhoods are color coded based on HR, p-value determined with a Cox proportional hazards model, and cluster stability across 5 iterations. **B)** Protein enrichment of GNP and GPP cell subsets determined using absolute Marker Enrichment Modeling (MEM). **C)** Overall survival of GNP-High (red line) and GNP-Low (black line) patients. **D)** Overall survival of GPP-High (blue line) and GPP-Low (black line) patients. **E)** The leftmost portion of the plot shows abundance of GNP and GPP cells identified by VR-Claw across 5 folds of identification on training data. GNP abundance is shown in red, and GPP abundance is shown in blue. GNP-High and GPP-High abundance thresholds are shown with red and blue dotted lines, respectively. The two columns in the middle of the plot show patient status (GNP-High in red, GPP-High in blue, and Low for both populations in grey) as determined by Velociraptor (VR, middle left column) and RAPID (R, middle right column). The rightmost portion of the plot shows overall patient survival. The dotted black line indicates a median overall survival of 389 days. Censored patients are indicated with *. **F)** VR-Eye analysis seeking Population 1 using the reference MEM label shown in (B). A spectrum intensity scale indicates cell similarity with red indicating high similarity and purple indicating low similarity. **G)** VR-Eye analysis seeking Population 2 using the reference MEM label shown in (B). A spectrum intensity scale indicates cell similarity with red indicating high similarity and purple indicating low similarity. **H)** Correlation between each patient’s percentage of GNP-like cells identified in Training Data and Testing Data by VR-Eye in the first round of 5-fold cross-validation. Pearson correlation tests were conducted. **I)** Correlation between each patient’s percentage of GPP-like cells identified in Training Data and Testing Data by VR-Eye in the first round of 5-fold cross-validation. Pearson correlation tests were conducted.

Across five folds of cross validation, patients were consistently classified as being either GNP-High or GPP-High based on their abundance of these prognostic subsets (**Figure 2C** and **Supplementary Figure S1**). Notably, patients with a large abundance of either GNP or GPP cells lacked the other prognostic subset (**Figure 2C**). Kaplan-Meier analyses confirmed that GNP-High patients have a shorter overall survival time and that GPP-High patients have a longer overall survival time compared to patients that had a low abundance of these two populations, respectively (**Figure 2D** and **Figure 2E**). VR-Claw accurately classified patients as being GNP-High or GPP-High compared to the previous classification reported by RAPID (GNP-High F1-measure=0.94, GPP-High F1- measure=0.72; **Supplementary Figure S1C**). These results indicated that VR-Claw successfully identified prognostic populations of GBM cells in **<u>Dataset 1</u>**.

#### Velociraptor-Eye accurately identified known populations of GBM cells based on learned phenotype

The next analysis goal was to establish that the novel cell seeking approach used by VR-Eye accurately identified cell populations based solely on their phenotype. MEM labels learned by VR- Claw were used as reference phenotypes for VR-Eye to seek in both the training data that was used to learn the phenotype and in the testing data that was withheld from VR-Claw and MEM analyses (**Figure 2F** and **Figure 2G**). Patient-level abundance of GNP-like cells by VR-Eye in the training data and testing data was highly correlated (R^2^=0.98; **Figure 2H**). Patient-level abundance of GPP-like cells by VR-Eye in the training data and testing data was highly correlated (R^2^=0.99; **Figure 2I**).

To determine the cellular features that best distinguished each population from other cells, a series of VR-Eye searches was run using different combinations of markers to compare cell identity to the population of interest. Populations within **<u>Dataset 1</u>** were used as “knowns” to be sought based on their multidimensional patterns of protein expression **Supplementary Figure S2A**. Specifically, RAPID-identified GNP and GPP populations were used to test VR-Eye. F1-measures were calculated to compare VR-Eye cell classification with RAPID cell classification. Optimization of search input revealed that the key cellular features that distinguished GNP cells from other cells were co-expression of SOX2 and S100B as well as lack of GFAP and p-NFkB expression (F1-measure=0.80; optimized GNP label: SOX2^+7^ S100B^+4^ GFAP^+1^ p-NFkB^+0^; **Supplementary Figure S2B**). GPP cells were distinguished by high expression of EGFR and lack of CD49F, p-p38, CD44, and B3TUB (F1- measure=0.90; optimized GPP label: EGFR^+7^ CD49F^+2^ p-p38^+2^ CD44^+1^ B3TUB^+0^. Population abundances for GNP and GPP cells identified by VR-Eye with the optimized search input were strongly correlated with population frequencies identified by RAPID (**Supplementary Figure S2C**; GNP R^2^=0.94, GPP R^2^=0.99).

To benchmark Velociraptor performance with other methods of population identification, VR-Eye and RAPID were each applied to ten different t-SNEs generated using the same sample of 131,880 GBM cells. Additionally, four cytometry experts manually gated for GNP and GPP cells based on a published gating scheme ^14^. VR-Eye identified GNP-like cells with a median F1-measure of 0.77 and GPP-like cells with a median F1-measure of 0.84 across ten runs (**Supplementary Figure S2D**). RAPID identified GNP-like cells with a median F1-measure of 0.63 and GPP-like cells with a median F1-measure of 0.61 across ten runs. Biaxial gating identified GNP-like cells with a median F1- measure of 0.35 and GPP-like cells with a median F1-measure of 0.57 across ten runs. Taken together, these analyses established that VR-Eye accurately and reproducibly identified cell populations by seeking a specified phenotype.

#### Velociraptor-Eye accurately revealed cell populations in individual analyses

The next goal was to compare a multi-sample pooled analysis to individual sample analyses. In this test of VR-Eye, all GBM cells from each tumor in **<u>Dataset 1</u>** were analyzed in a patient-specific manner rather than in a down-sampled, combined analysis. This type of analysis also determines how representative a subset of cells is to an entire tumor sample. For each tumor, all GBM cells were used to generate a tumor-specific t-SNE (**Figure 3A**). The number of GBM cells in individual tumors ranged from 4,710 cells to 329,650 cells with a median cell count of 68,761. Next, GNP and GPP populations were sought with VR-Eye using the optimized population phenotypes (**Figure 3B**). The majority of GBM cells were not similar to either prognostic phenotype. Individual tumors ranged in GNP-like abundance from 0.00% to 54% and in GPP-like abundance from 0.00% to 88%. Tumors with greater than 5% abundance either GNP-like or GPP-like cells contained less than 2.5% of the other population. The difference between each cell’s GNP similarity score and GPP similarity score was calculated for each sample as a surrogate of population homogeneity (**Figure 3C**). Population abundances for GNP and GPP cells identified by VR-Eye in full tumor samples were strongly correlated with population frequencies identified by RAPID in the down-sampled dataset (**Figure 3D**; GNP R^2^=0.96, GPP R^2^=0.98). These results show that VR-Eye accurately identifies cell populations in individual samples as opposed to a pooled analysis.

**Figure 3.**
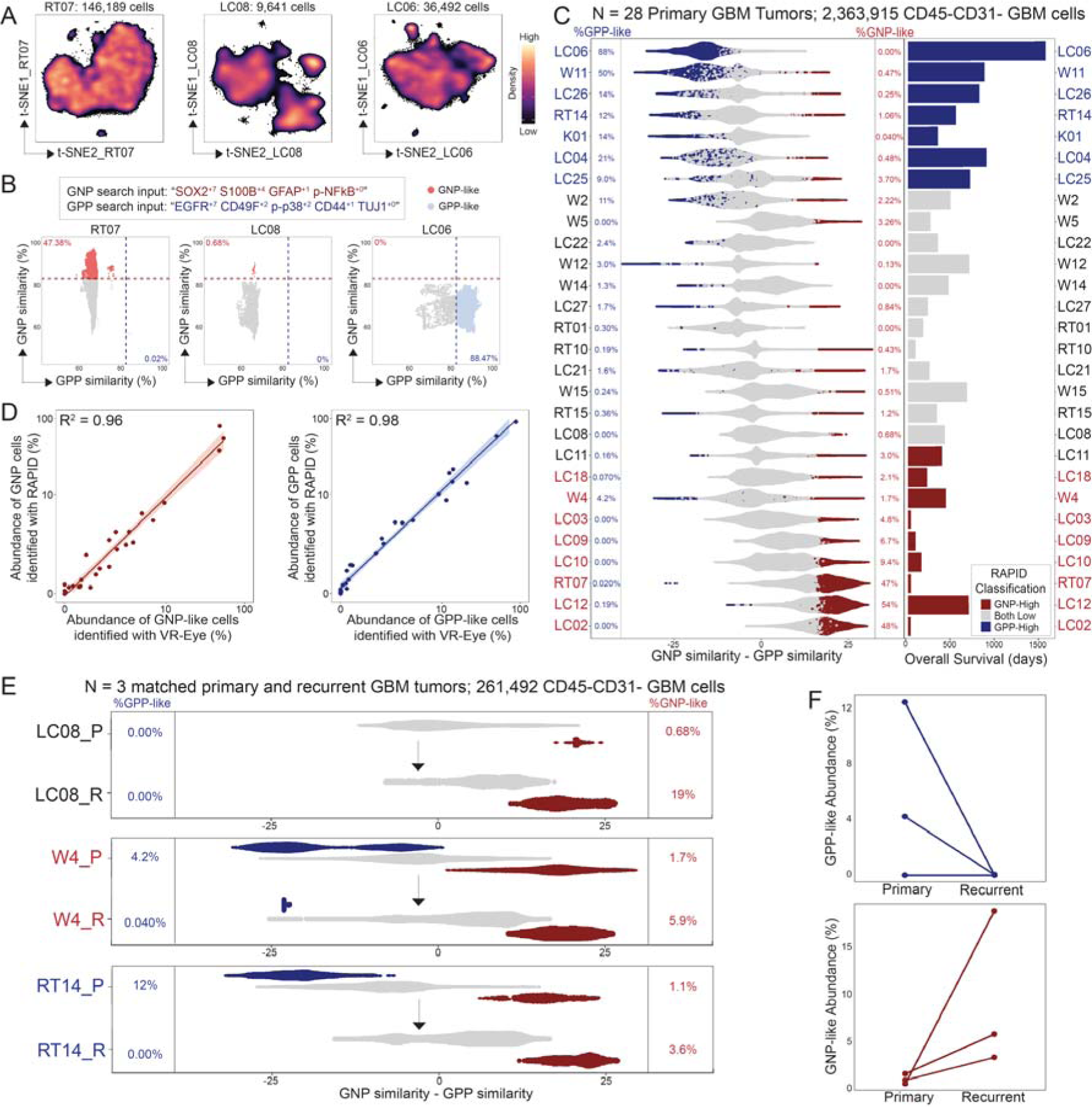
Velociraptor-Eye quantifies opposing cell identities and reveals shifts in glioblastoma tumor cell composition at recurrence. **A)** Three representative t-SNEs each created using a single patient’s CD45^-^CD31^-^ GBM cells (individual cell counts ranged from 4,710 to 329,650 per patient for a total of 2,363,915 live GBM cells). A magma intensity scale represents cell density on the t-SNE axes. **B)** Three representative VR-Eye analyses of individual, patient-specific t-SNEs generated from CD45^-^CD31^-^ GBM cell populations. Plots display each patient’s cells’ similarity to GNP cells on the y-axis (GNP searching label: SOX2^+7^ S100B^+4^ GFAP^+1^ p-NFkB^+0^) and GPP cells on the x-axis (GPP searching label: EGFR^+7^ p-p38^+2^ CD49F^+2^ CD44^+1^ TUJ1^+0^). The dotted red horizontal line indicates the GNP-like similarity threshold of 83%, such that every cell above this line is considered “GNP-like”. The dotted blue vertical line indicates the GPP-like similarity threshold of 82%, such that every cell to the right of this line is considered “GPP-like”. **C)** Patient-level VR-Eye summary. Red dots represent GNP-like cells, and blue dots represent GPP-like cells as determined by VR-Eye. Patients are ordered by decreasing range of %GPP cells to %GNP cells as determined by RAPID. Bars to the right represent overall survival time. Patient codes are colored based on RAPID status, where red indicates a GNP-High patient and blue represents a GPP-High patient according to RAPID. **D)** Correlation between each patient’s percentage of GNP-like or GPP- like cells, respectively, identified by RAPID in Dataset 1 and by VR-Eye on the entire GBM cell population per tumor. **E)** VR-Eye analyses of individual patient t-SNEs generated from CD45^-^CD31^-^ GBM cell populations from three matched primary and recurrent GBM (individual t-SNEs range from 7,498 to 134,876 cells per patient). Density plots show the difference between each cell’s GNP and GPP similarity. Cells colored blue had a GPP similarity value greater than 82% and are considered “GPP-like”. Cells colored red had a GNP similarity value greater than 83% and are considered “GNP-like”. Primary tumors are labeled with “_P”, and recurrent tumors are labeled with “_R”. **F)** Quantification of GNP-like and GPP-like cell abundance in paired primary and recurrent tumors.

### Velociraptor-Eye identified prognostic cell populations in recurrent GBM tumors

Having verified that VR accurately and reproducibly identified prognostic cell populations that have been reported in a published dataset, the goal of the next analysis was to use Velociraptor to identify prognostic cells in newly collected data. To address this goal, three recurrent GBM tumors were resected, dissociated, and stained with a 33-dimensional antibody panel to measure protein expression using mass cytometry as previously described (**<u>Dataset 2</u>**) ^14,29^. All 33 markers in the antibody panel used to characterize the recurrent tumors were present and measured in the same channels in **<u>Dataset 1</u>**, and each recurrent tumor had a corresponding primary tumor that was included in **<u>Dataset 1</u>**. For each recurrent tumor, VR-Eye sought GNP and GPP populations using the optimized phenotypes for each population, respectively (**Figure 3E**). Each recurrent tumor contained a greater percentage of GNP-like cells than its corresponding primary tumor, and in the two primary tumors that originally contained GPP-like cells, the corresponding recurrent tumors each showed a decrease in GPP-like cell abundance (**Figure 3F**).

### Velociraptor identified genotype-specific cells in patients with Inborn Errors of Immunity

The next test of Velociraptor was designed to assess how well the entire workflow performs in the case of limited sample numbers and limited cell numbers. To address this question, we performed deep T cell profiling on PBMCs from six sets of patients with Inborn Errors of Immunity and healthy donors using a 43-dimensional mass cytometry panel (**<u>Dataset 3</u>**). Inborn Errors of Immunity encompass a variety of rare, monogenic mutations that lead to diverse clinical presentations of autoimmunity, autoinflammation and immunodeficiencies ^30,31^. The first set of donors included data from 1 healthy adult donor and 2 IEI patients with STAT1 GOF mutations that have been previously reported ^32^ as well as unpublished data from 2 patients with *GATA2* haploinsufficiency (Set 1, N=5). Donor sets 2-6 included an additional 38 IEI patients, 6 healthy adult donors, and 2 healthy pediatric donors (N=46). In total, all donors included in this study (N=51) represented mutations in 26 unique genes. The inclusion of multiple patients with *GATA2* haploinsufficiency in two batches that were collected on different dates presented the opportunity to identify *GATA2* haploinsufficiency-specific cell populations in Set 1 that could then be sought in Set 2 for validation.

T cells from donors in Set 1 were embedded into a two-dimensional t-SNE based on protein expression (**Figure 4A**). A modified version of VR-Claw was created to identify cells associated with a categorical variable (e.g., genotype) using local phenotypic neighborhood-based identification. This version of VR-Claw was inspired by T-REX ^10^, but differs in two key aspects: 1) VR-Claw tests each category represented in a dataset for cell enrichment rather than being restricted to a binary comparison, and 2) VR-Claw includes a series of filtering steps that ensure findings are not specific to a single patient within a cohort analysis. VR-Claw identified three populations that were greatly enriched in (>95%) and three populations that were greatly lacking from (<5%) the patients with *GATA2* haploinsufficiency (**Figure 4B**). Further inspection of these populations confirmed that populations were consistent in both *GATA2* haploinsufficient patients (**Figure 4C**). MEM was used to quantify the phenotypes of each cell population (**Figure 4D**). CD4^+^ and CD8^+^ T cells populations that were statistically lacking from the two patients with *GATA2* haploinsufficiency each expressed high levels of CD27 and CCR7 (MEM scores ≥ +6). Conversely, the CD4^+^ and CD8^+^ populations that were statistically enriched in the two patients with *GATA2* haploinsufficiency expressed very high levels of CD57 (MEM scores ≥ +6). VR-Claw was repeated an additional nine times using independently sampled cells with replacement to verify population stability (**Supplementary Figure S3**).

**Figure 4.**
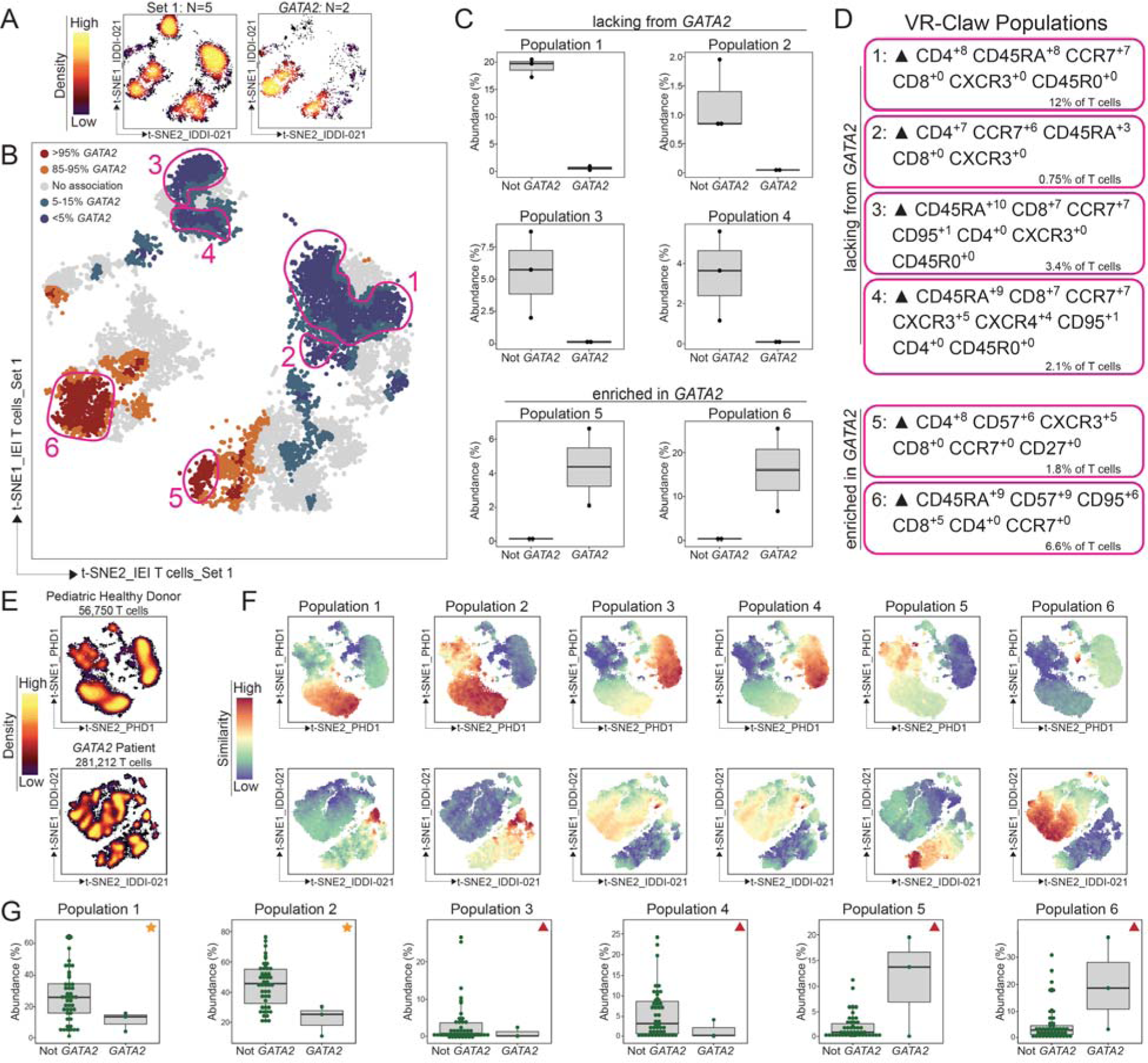
Velociraptor reveals, characterizes, and confirms CD57^+^ T cells that are enriched in patients with *GATA2* haploinsufficiency. **A)** Cell density on t-SNE axes generated using T cells from donors in Set 1. Set 1 includes one adult healthy donor, two IEI donors with *GATA2* haploinsufficiency, and two IEI donors without *GATA2* haploinsufficiency. The left plot shows equally sampled T cells from all patients in Set 1 (N=5 donors, n=10,000 cells). The right plot shows equally sampled T cell from the *GATA2* haploinsufficiency patients in Set 1 only (N=2 donors, n=4,000 cells). A magma color scale indicates cell density on the t-SNE axes where purple indicates low density and yellow represents high density. **B)** VR-Claw analysis on equally sampled T cells from IEI patients and healthy donors in Set 1 (N=5 donors, n=10,000 cells). Red and orange denote populations enriched in *GATA2* haploinsufficiency patients; blue and purple denote populations lacking from *GATA2* haploinsufficiency patients. **C)** Quantification of cell abundance with respect to all T cells for each VR-Claw population. **D)** Filtered MEM labels show proteins enriched in VR-Claw Populations. Proteins with a relative MEM score of > 2 were included. **E)** Cell density on t-SNE axes. The top plot was generated using all T cells from a healthy pediatric donor included in Set 2 (n= 55,201 cells). The bottom plot was generated using all T cells from an IEI patient with *GATA2* haploinsufficiency included in Set 1 (n= 278,870 cells). A magma color scale indicates cell density on the t-SNE axes where purple indicates low density and yellow represents high density. **F)** Representative VR-Eye analyses seeking VR-Claw Populations 1- 6 in T cells from a pediatric healthy donor (PHD1, top row) and a *GATA2* haploinsufficient patient (IDDI-040, bottom row). MEM labels shown in Panel D were used as the searching label for each population. A rainbow intensity scale indicates similarity to the sought population with red representing high and purple representing low similarity. **G)** Quantification of T cell population abundances in all IEI patients and healthy donors (N=51) as determined by VR-Eye. Gold stars indicate high priority populations to continue studying, and red triangles indicate populations that require further validation using additional *GATA2* haploinsufficiency samples.

VR-Eye was then used to seek similar cell populations in donor samples from Sets 2-6. All T cells from each donor were embedded into donor-specific t-SNE spaces to maximize the number of cells included in this analysis (**Figure 4E**). Each population revealed with VR-Claw was first sought in samples from Set 1 to validate findings and to determine appropriate similarity thresholds that distinguish each population (**Figure 4F**). Each population was then sought in the remaining IEI patient and healthy donor samples (**Figure 4F**). For each patient, the abundance of each population was quantified (**Figure 4G**). Taken together, these analyses showed that Velociraptor successfully identified clinically relevant cells in the case of limited samples and limited cell numbers.

### Cross-platform identification

Having shown that VR-Eye accurately identifies cells that were characterized using mass cytometry, the goal of the next analysis was to establish whether VR-Eye could be used to identify cells characterized with other single cell platforms. This question was addressed using a 41-dimensional immune-focused imaging mass cytometry dataset that characterized tumor immune microenvironments of human breast cancer (**<u>Dataset 4</u>**) ^33^. In this dataset, the authors annotated 18 different cell types. To first establish that VR-Eye could accurately identify cells in a new data type, each cell type identified by Tietscher et al. was sought and compared to the original identification (**Figure 5A**). Each population was sought using a 41-feature phenotypic quantification that included every protein measured in the dataset as well as with a filtered phenotypic quantification that only included proteins that were specifically enriched or lacking in that population according to relative MEM on a common t-SNE embedded with equally sampled cells from each patient (n=257,076 cells total; **Supplementary Figure S4A**). Filtered search input ranged in size from 4 features to 28 features. Full 41-feature labels resulted in more accurate identification than the filtered labels in seven out of ten cell types (**Supplementary Figure S4B**). To compare the populations identified by the original authors and by VR-Eye, a phenotypic homogeneity score was calculated. This score compared how similar a single neighborhood’s phenotype was to the phenotype of its corresponding population. Populations identified by VR-Eye using full 41-dimensional labels had the highest homogeneity scores in nine out of ten cell types compared to the original populations and the populations identified by VR-Eye with a filtered label (**Supplementary Figure S4C**). Cell identification accuracy was confirmed using LOOCV. These results showed that VR-Eye accurately identifies cell populations in an external imaging mass cytometry dataset.

**Figure 5.**
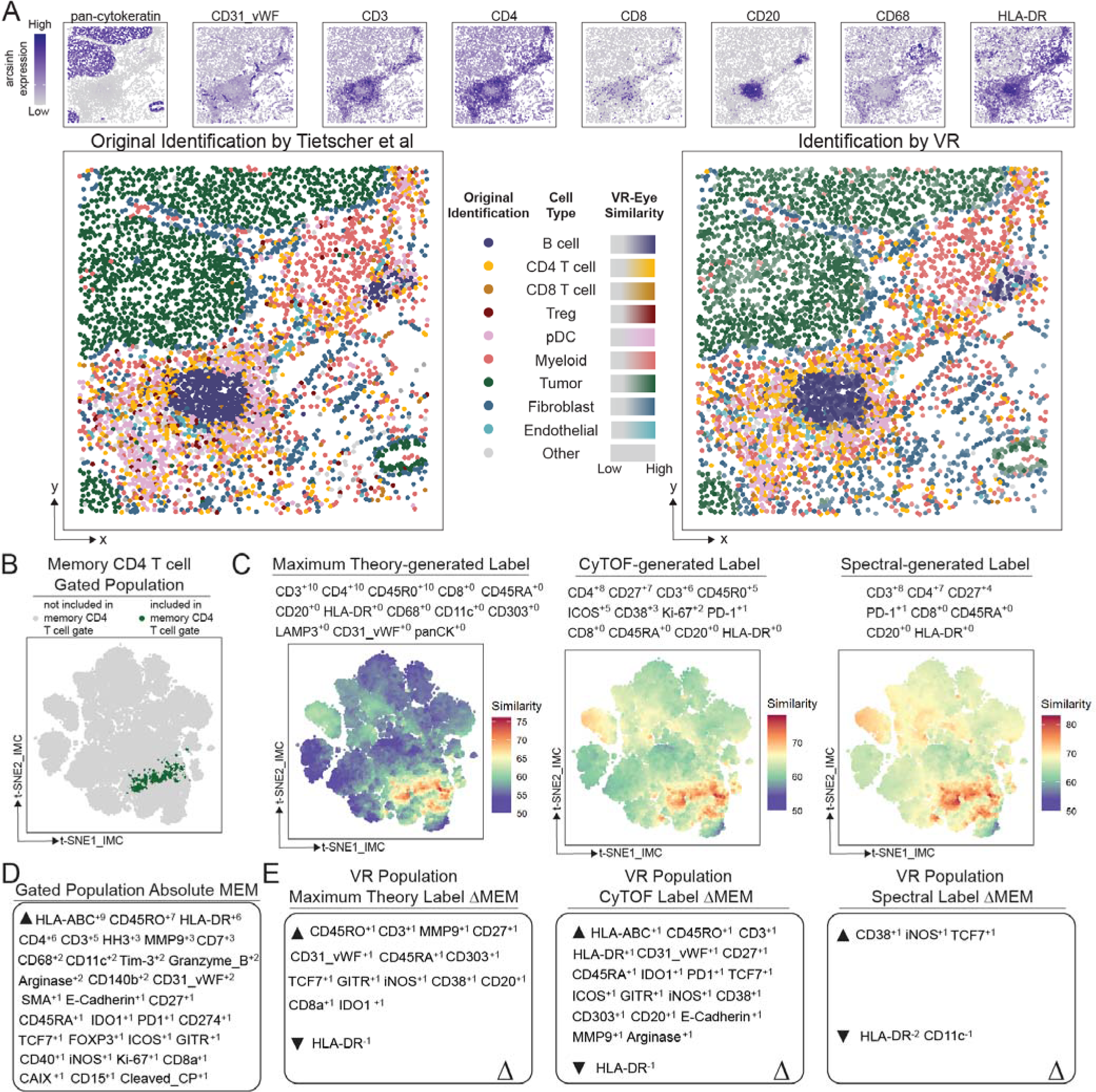
VR-Eye accurately identifies cell types in an external imaging dataset using reference phenotypes generated by different single-cell platforms. **A)** Plots along the top of the panel show the expression of selected proteins on an arcsinh scale. Grey indicates low expression and purple indicates high expression. The bottom left plot shows cell types as determined by Tietscher et al on a representative slide. The bottom right plot shows VR-Eye analyses for each cell type shown in the upper left plot. VR similarity was plotted as a gradient for each cell type as indicated. **B)** All IMC cells (n=257,076) embedded in a common t-SNE. Cells identified as memory CD4 T cells by biaxial gating are plotted in dark green. **C)** VR-Eye analyses seeking memory CD4 T cells based on reference populations measured with various platforms. The theory-generated label was written based on known protein expression patterns of memory CD4 T cells. The CyTOF-generated label was calculated on memory CD4 T cells that were gated in a healthy donor from Dataset 3. The spectral-generated label was calculated on memory CD4 T cells that were gated in Dataset 5. A rainbow intensity scale indicates similarity to the sought population with red representing high and purple representing low similarity. **D)** MEM label generated from the gated memory CD4 population. **E)** ΔMEM analyses comparing MEM labels from each VR-identified memory CD4 T cell populations to the MEM label generated from the gated memory CD4 T cell population.

To test whether quantified reference populations can be used to identify similar cells across data types, we sought memory T cells using a variety of search inputs that were generated using different technologies. Memory CD4 and CD8 T cells subsets were manually gated as a reference population (**Supplementary Figure S5A**). Theoretical memory T cell labels were generated based on known patterns of protein expression. Minimal theoretical labels including CD3, CD45R0, and CD4 or CD8a markers were designed as the minimal feature set that could be used to identify subsets of memory T cells, and maximum theoretical labels including CD3, CD4, CD8a, CD45R0, CD45RA, CD20, HLA-DR, CD68, CD11c, CD303, LAMP3, CD31_vWF, and panCK markers were designed to exclude additional cell types present in the tissue. To assess how well reference phenotypes calculated from orthogonal platforms could identify memory T cells, MEM was performed on a biaxially gated memory T cell populations from a healthy adult donor in **<u>Dataset 3</u>** to yield a CyTOF-generated label, and a 40-dimensional spectral flow cytometry dataset (**<u>Dataset 5</u>**) ^34^ was used to calculate a spectral- generated MEM label on memory T cell populations. Each of these labels were used to seek memory T cells with VR-Eye (**Figure 5B**, **Figure 5C, and Supplementary Figure S5B**).

Memory CD4 and CD8 subsets were identified with VR-Eye using similarity thresholds that were optimized to best capture the gated population. The phenotype of each VR-identified population was compared to biaxially gated memory T cell populations with a ΔMEM analysis (**Figure 5D**, **Figure 5E, and Supplementary Figure S5C**). Features of VR populations identified using the maximum theory- generated label and the CyTOF-generated label did not differ from the MEM label of the gated memory CD4 T cell population by more than 1. This indicates high phenotypic similarity between the VR-identified population and the gated population. The greatest difference observed was elevated CD8a expression on the VR population identified using the minimum theory-generated label (ΔMEM: CD8a^+2^). To compare the VR-identified populations to the gated populations at the individual cell level, the precision, recall, and F1-measure for each VR-Eye analysis was calculated (**Supplementary Figure S5D**).

Upon inspection of the data, we observed cells that shared high similarity to both memory CD4 and CD8 T cell populations in addition to cells that strongly matched a single phenotype, and these cells were confirmed with a VR-Eye search for CD4^+^CD8^+^ double positive memory T cells (**Supplementary Figure S5E**). The difference between each cell’s memory CD8 similarity score and memory CD4 similarity score was calculated to distinguish cells that matched only a single phenotype (**Supplementary Figure S5F**). To exclude cells that matched multiple memory T cell subsets, we filtered the VR-identified memory CD4 T cell population such that each cell (1) surpassed the optimized memory CD4 T cell similarity threshold, and (2) had a difference in memory CD4 and memory CD8 similarity of greater than 5. This filtering step increased the percentage of VR-identified memory CD4 T cells falling into the CD4^+^ biaxial gate from 66% to 90% (**Supplementary Figure S5G**). Exclusion of cells that matched both memory CD4 and CD8 phenotypes also improved population recall compared to gated populations (memory CD4 recall = 0.93, memory CD8 recall = 0.97). These results show that VR-Eye enables cross-platform cell identification and accurately identifies sought populations using different feature sets.

## Discussion

Velociraptor identifies cells based on association with external variables or similarity to a reference phenotype. The modular nature of the workflow follows best practices developed for cytometry analysis in prior decades ^35,36^ and allows for independent use of a single tool or a complete analysis using VR-Claw and VR-Eye linked by MEM. The use of local phenotypic neighborhoods that track across continuous phenotypic space rather than placing each cell into a single cluster allows Velociraptor to identify both abundant and rare populations of cells at a granular level and to discern subtle changes in protein expression across populations. VR-Claw identified expected GNP-like and GPP-like populations in **<u>Dataset 1</u>** that were identified in previous studies using the RAPID algorithm^14^ with exact and reproducible boundaries around prognostic clusters that were significantly associated with patient outcome (p<0.01; **Figure 2A**). Notably, the local neighborhood approach enabled VR-Claw to identify specific regions of GPP-like cells more significantly associated with outcome (p<0.01, HR<1, [95% CI 0.039-0.493]) compared to those identified with RAPID, as well as an additional GPP cluster (cluster 3) that co-expresses S100B and EGFR and was not found by RAPID due to limitations in RAPID’s clustering approach. Therefore, VR-Claw can be used to reveal rare prognostic cells that have historically been overlooked.

VR-Eye offers a novel approach to defining cell identity that is based on phenotypic comparisons with well-established cell identities to assess if a population strongly matches a single cell type or if it has similarity with multiple populations. Additionally, VR-Eye simultaneously considers an entire phenotype when calculating a continuous similarity score rather than performing sequential binary filtering based on strict thresholds that distinguish marker negativity and positivity; this allows VR-Eye to tolerate a small degree of variability in the expression of a protein. These features of VR-Eye allow for multidimensional definitions of cell identity rather than binary cell classification. VR-Eye outperformed RAPID and biaxial gating cell identification strategies by reproducibly identifying the same GNP and GPP cells across 10 t-SNEs generated using the same sample of 131,880 GBM cells (**Supplementary Figure S2**). VR-Eye also enables cell identification within an individual sample rather than within a pooled analysis, which bypasses the need to re-analyze historical cohorts upon collection of a new sample. Additionally, VR-Eye interrogates every cell in a sample rather than requiring equal sampling based on the sample with the fewest cells in a grouped analysis, which increases the total number of cells included in an analysis and thus the overall power of the analysis.

Notably, Velociraptor revealed that GBM cell populations associated with shorter survival times were more abundant in GBM tumor recurrences compared to the same patient’s primary GBM tumor (**Figure 4E** and **Figure 4F**). Conversely, populations of positive prognostic GBM cells were greatly diminished or completely lacking in matched recurrent tumors. These shifts in population frequency occurred regardless of the original frequency of GNP and GPP cells in the primary tumor. One potential explanation for this change in tumor makeup is that GPP cells are more susceptible to chemotherapy and radiation treatment, whereas GNP cells may persist such that the same GNP cells and their progeny remain in the recurrent tumor. An alternative explanation is that another population of cells survives treatment and later gives rise to a new population of GNP cells in the recurrent tumor either due to plasticity or differentiation. Interestingly, the phenotype of GNP cells share similarity with multiple healthy neuronal cells due to aberrant co-expression of S100B, a marker of astrocytic cells, and SOX2, a Yamanaka factor that is highly expressed in pluripotent stem cells ^1,37^. Due to the increased expression of the transcription factor SOX2, GNP cells can potentially be considered more “stem-like” than their GPP counterparts. This feature of cell identity may contribute to their persistence and increased abundance in recurrent GBM tumors compared to matched primary GBM tumors, as cancer stem cells have been implicated as major players driving tumors and tumor recurrence across cancer types ^38–41^.

As a proof of concept, Velociraptor next identified genotype-specific cells across sets of patients with IEIs. VR-Claw first revealed multiple CD57^+^ T cell subsets that were statistically enriched in IEI patients using samples of only 2000 cells per patient. VR-Eye then sought similar cell populations in other healthy donors and IEI patients, including one additional patient with *GATA2* haploinsufficiency. Results suggested a lack of naïve T cell subsets in IEI patients with *GATA2* haploinsufficiency and a corresponding enrichment of CD57 T cell subsets; however, additional samples are required to confirm these findings. Notably, CD57 cells have been observed in other disease settings. After observing these findings from Velociraptor, we noted a study that also reported that patients with *GATA2* haploinsufficiency have reduced numbers of naïve T cells and increased numbers of CD57^+^ T cells compared to healthy donors that correlated with clinical severity ^42^. This study supports the Velociraptor findings shown here, which confirms the usefulness of this workflow even in the case of limited patient samples (N = 3 patients with *GATA2* haploinsufficiency) and limited cell numbers (n = 2000 cells per patient in the VR-Claw analysis).

Finally, we used VR-Eye to identify cells across single cell platforms. Reference population labels generated using different platforms successfully identified memory CD4 T cells in the example shown here. Interestingly, a minimum theoretical phenotypic description that contained only markers thought to be highly expressed in the sought population did not accurately identify memory CD4 T cells. Instead, searching labels containing a balance of features that are highly expressed (+10) and not expressed (+0) performed well across datasets and single cell modalities. This feature of a robust searching label was also observed in the optimization of reference phenotypes in Dataset 1 and Dataset 3, suggesting that cell identity can be described in terms of both positive and negative enrichment. Reference phenotypes generated using immune-focused panels from CyTOF and spectral flow cytometry successfully identified memory CD4 T cells despite only sharing 12 and 8 markers, respectively, with the IMC antibody panel. Inclusion of additional markers, such as panCK to discriminate against malignant cells, in the maximum theoretical label did increase the difference between identified memory CD4 T cells and distinct cell types. This analysis also revealed a small population of CD4^+^ cells that had been excluded from the CD4 T cell memory compartment with expert gating. Upon manual inspection, these cells closely matched the memory CD4 label, but also expressed CD8a which was likely why they were excluded from the gated population. A subsequent search found these cells matched a CD4^+^CD8^+^ double positive memory label greater than a CD4^+^ single positive label. Overall, these results (1) helped resolve a population that had not been described in the original research or by manual gating, and (2) demonstrated the value of an objective match, of including both positive and negative features, and of detecting matches to more than one known identity.

Additional algorithms have been developed by others to integrate data collected by multiple single cell modalities by placing each cell into a common phenotypic space following data transformation, regardless of the origin of the data. However, this method may require many shared features between datasets as it was developed for single cell transcriptomics ^22,43^ and relies on the presence of similar cell types and cell states in each dataset ^24^. VR-Eye differs in that it does not require *a priori* knowledge that similar cell populations are present in two datasets, but rather tests if there are similar cells present in two datatypes. Through utilization of MEM, orthogonal data types need not be forced into common phenotypic space. Thus, VR-Eye can be used to seek similar populations of cells across data types, disease states, and types of tissue.

Both tools in the Velociraptor workflow perform phenotypic neighborhood definition on dimensionally reduced latent spaces. Dimensionality reduction preserves high-dimensional data structure while condensing data into a space that is less sparse and more amenable to clustering ^44^. One type of dimensionality reduction that was not compared here is a graphical representation of the data ^12^. One benefit of using a graphical data structure is that many downstream analysis tools are designed to use graphical data structures as input ^45–47^. However, dimensionally reduced data spaces generated via t-SNE or UMAP can be converted to graph representation if so desired, where each cell is a node and edges are drawn between each cell and its phenotypic neighbors.

Although the Velociraptor workflow successfully identified clinically relevant cell populations across human disease datasets, there are some limitations to the approach. First and foremost, rigorous experimental design and execution remain paramount to accurate cell identification using unsupervised machine learning. As with pooled analysis, proper data QC and batch normalization improve results from VR-Claw. Batch effects are less of a concern when performing individual analyses using VR-Eye so long as the overall pattern of protein expression remains consistent across batches. Prior to data collection, careful consideration and design of antibody panels will determine how well Velociraptor can perform. A panel must have sufficient overlap in features with any dataset it is to be compared to, so that populations can be identified across datasets. Additionally, a MEM label is designed to capture the full range of data that can be collected on a given single cell platform. Performing MEM on a small subset of cells that does not represent the entire data scale of a platform may inflate the enrichment levels of proteins which could lead to inaccurate downstream cross-dataset identification. VR-Eye uses a reference phenotype to find similar cells across datasets, and the specification of features in the searching label is the most crucial step when using this algorithm. Inclusion of too many features may be over-optimized to the specific set of cells used to generate that label. We therefore recommend careful optimization and validation of a search label to ensure its generalizability. Standard statistical validation approaches used here, including cluster stability testing and cross validation, should be performed to confirm any Velociraptor findings.

With the rapid increase in high-dimensional single cell technologies that can generate distinct information about cell populations, there is a need to match similar cells across datasets to learn a more detailed understanding of cell identity. Therefore, robust measurements of cell identity are required to accurately identify cells across data types and thus to hypothesize about their role in disease. Multiple continuous cell comparisons to well-established populations using Velociraptor will provide a new perspective on cell identity that could be used to implicate similar cell populations across disease settings.

## Methods

### GBM Tissue Collection and Processing for Dataset 2

Recurrent GBM tumors were surgically resected at Vanderbilt University Medical Center between 2014 and 2017. All samples were collected with written informed consent under Institutional Review Board protocol #131870 and in accordance with the Declaration of Helsinki. Tumor samples were dissociated into single cell suspensions as previously described ^29^.

### IEI and Donor PBMC Collection for Dataset 3

Blood was collected from IEI patients with written informed consent under Institutional Review Board protocol #182228, from healthy donors with written informed consent under Institutional Review Board protocols #131311 and #191562, and in accordance with the Declaration of Helsinki. 100mL of blood per donor were collected via venipuncture into heparin tubes (Becton Dickinson). Blood was diluted 1:4 with PBS, placed into a Ficoll-Paque Plus density gradient (GE Lifesciences), and centrifuged at 400 x g for 30 min. Next, buffy coats were isolated, washed with PBS, and centrifuged at 500 x g for 10 min. Cell pellets were resuspended in ACK lysis buffer for 5 min, washed with X, and cryopreserved at 1 x 10^7^ cells/mL in 10% DMSO in FBS at 80°C.

### Metal-conjugated antibodies

All antibodies used for mass cytometry analysis are listed in **Supplementary Table 1 and Supplementary Table 2**. Pre-conjugated, metal-tagged antibodies were purchased from Fluidigm, and unconjugated pre-conjugated purchased in purified form and custom conjugated using the MaxparX8 Antibody Labeling Kit (Fluidigm) according to manufacturer’s protocol.

### Cell Preparation and Mass Cytometry

Antibody staining and mass cytometry analyses were performed as previously described ^48,49^. Cryopreserved samples were rapidly thawed in a 37°C water bath and resuspended in complete RPMI 1640 supplemented with 10% FBS and 50 U/ml penicillin-streptomycin (Thermo Scientific HyClone). Cells were washed with serum-free RPMI 1640 and rested for 15 min. Next, cells were washed with serum-free RPMI 1640 and stained with a 103Rh Cell-ID intercalator (Fluidigm) at a final concentration of 1 μM for 5 min at RT. Cells were then washed with PBS + 1% BSA to quench staining. Samples were then stained with the appropriate surface antibody cocktail (**Supplementary Table 1 and Supplementary Table 2**) at RT for 30 min. Next, cells were washed with PBS and fixed with 2% formaldehyde at room temperature for 20 min. Cells were washed with PBS and permeabilized with ice cold methanol overnight at 20°C. The following morning, cells were washed with PBS then washed with PBS + 1% BSA. Next, samples were stained with intracellular antibodies at RT for 30 min and washed with PBS. Cells were intercalated with IridiumCell-ID at a final concentration of 125 nM in PBS + 1.6% formaldehyde at 4°C overnight. On the day of data collection, cells were washed once with PBS and washed once in ultrapure deionized water. Samples were resuspended in ultrapure deionized water with 10% EQ four element calibration beads (Fluidigm) and filtered through a 40-mm FACS filter tube. Data were collected on a Helios CyTOF 3.0 (Fluidigm) and stored in FCS files. Instrument quality control and tuning processes were performed following the guidelines for the daily instrument operation.

### Data Processing

Raw mass cytometry files were normalized using the MATLAB bead normalization tool ^50^ prior to upload to the cloud-based analysis platform Cytobank ^51^. Data were arcsinh transformed with a cofactor of 5. Cell doublets were excluded using Gaussian parameters, beads were excluded, and intact live cells were selected for downstream analysis based on DNA content measured via Iridium and Rhodium intercalation. In **<u>Dataset 2</u>**, CD45^-^CD31^-^ GBM cells were selected for downstream analyses. In **<u>Dataset 3</u>**, CD45^+^ cells from all donors were plotted on a common t-SNE, and T cells were selected based on CD3 expression. Donors with at least 2,000 T cells were included in subsequent analyses.

### Velociraptor-Claw Algorithm

The Velociraptor-Claw algorithm includes dimensionality reduction, definition of local phenotypic neighborhoods, iterative testing of cell neighborhoods for association with an external variable, and phenotypic quantification of identified cells of interest. Here, dimensionality reduction was performed on using t-SNE either in R or on Cytobank with a perplexity of 60 and 10,000 iterations. Local phenotypic neighborhoods were defined as the k-nearest neighbors (KNN) for every cell in the low dimensional embedding of the dataset using the fast nearest neighbors (FNN) package in R. The value of k was defined to be the square root of the total number of cells included in the analysis ^52^. Within each neighborhood, patients are categorized as Low or High for that neighborhood based on their abundance of cells residing in that neighborhood compared to the interquartile range of patient cell abundance for that neighborhood as described ^14^. A univariate Cox proportional hazards model was used to investigate the association of each neighborhood with patient overall survival time. The index cell of each neighborhood was annotated with its neighborhood’s effect size (hazard ratio) and statistical significance (p-value). Cells are then clustered into prognostic populations using DBSCAN (from the dbscan package in R). A univariate Cox proportional hazards model is then performed on each prognostic population to ensure that it is still associated with overall survival following DBSCAN clustering. Prognostic populations can be filtered by the number of cells in that cluster or by the number of samples in that cluster. Finally, the phenotypes of prognostic populations are quantified using MEM ^17^.

The modified version of VR-Claw used to analyze **<u>Dataset 3</u>**identified cells that were enriched (>95%) or lacking (<5%) from categories of patients within a dataset (i.e., genotype) following cell neighborhood definition. Genotype-specific cells were clustered using DBSCAN and filtered such that final clusters contained at least 25 cells and were consistently enriched/lacking across all patients within that genotype class. All steps downstream of dimensionality reduction in both versions of VR- Claw are deterministic.

### Velociraptor-Eye Algorithm

The Velociraptor-Eye algorithm includes dimensionality reduction, definition of local phenotypic neighborhoods, and phenotypic similarity calculation of each neighborhood compared to a reference phenotype. A reference phenotype that is written in the form of a MEM label is supplied as input. As described above, dimensionality reduction was performed on using t-SNE either in R or on Cytobank with a perplexity of 60 and 10,000 iterations. Local phenotypic neighborhoods were defined as the k-nearest neighbors (KNN) for every cell in the low dimensional embedding of the dataset using the fast nearest neighbors (FNN) package in R. The value of k was defined to be 60. MEM was used to quantify the phenotype of each cell neighborhood, and the similarity of each neighborhood to the reference population was calculated using the root-mean-square deviation (RMSD) between neighborhood and reference MEM labels (**Equation 1**).

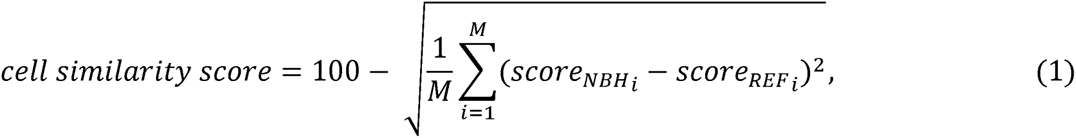

In Equation 1, M represents the number of markers shared between the reference MEM label and the MEM label of the cell neighborhood, score_NBH_ denotes the cell neighborhood’s MEM score for a particular marker, and score_REF_ denotes the reference label’s MEM score for the same marker. Following dimensionality reduction, each step of VR-Eye is deterministic.

## Data Availability

Datasets analyzed in this manuscript are online at FlowRepository ^53^, and will be made public upon acceptance. Transparent analysis scripts for datasets in this manuscript will be made available on the CytoLab Github page (https://github.com/cytolab/) with open-source code and commented Rmarkdown analysis walkthroughs upon acceptance.

## Acknowledgements

We thank Vanderbilt’s Cancer and Immunology Core as well as all the surgeons, patients, and families that supported this work. We thank Human Immunology Discovery Initiative collaborators, including Todd Bartkowiak, for helpful discussions of the inborn errors of immunity data, and we thank Caroline Roe for discussions of cell identification algorithms. Research was supported by the following funding resources: NIH/NCI grants R01 NS096238 (RAI, JMI), R01 CA226833 (JMI, CEC, SM, MJH), R01 NS118580 (RAI), U01 AI125056 (JMI), U54 CA217450 (JMI, MJH), T32GM137793 (CEC), the Vanderbilt-Ingram Cancer Center (VICC, P30 CA68485), the Michael David Greene Brain Cancer Fund (RAI, JMI), the Southeastern Brain Tumor Foundation (RAI, JMI), a gift from Daniel F Hewins (RAI), the Ben & Catherine Ivy Foundation (RAI, JMI), and by the Human Immunology Discovery Initiative of the Vanderbilt Center for Immunobiology.

## Author Contributions

CEC and JMI designed the study and conceptualized the velociraptor workflow. MJH and SM collected data. CEC and CM coded data analysis scripts. CEC, CM, and JMI performed mass cytometry data analysis and interpretation. SK and JAC provided clinical care to patients with IEIs and identified GATA cases. RAI, JMI, LBC, and RCT developed Vanderbilt’s human glioblastoma research program. RAI, MJH, and SM coordinated intraoperative tissue collection by research teams. LBC, RCT, and KDW provided freshly resected glioblastoma tissue specimens, including recurrence samples. CEC and JMI wrote the manuscript. JCR, RAI, and JMI provided financial support. All authors contributed to reviewing and editing the manuscript.

---

**Supplementary Figure S1.**
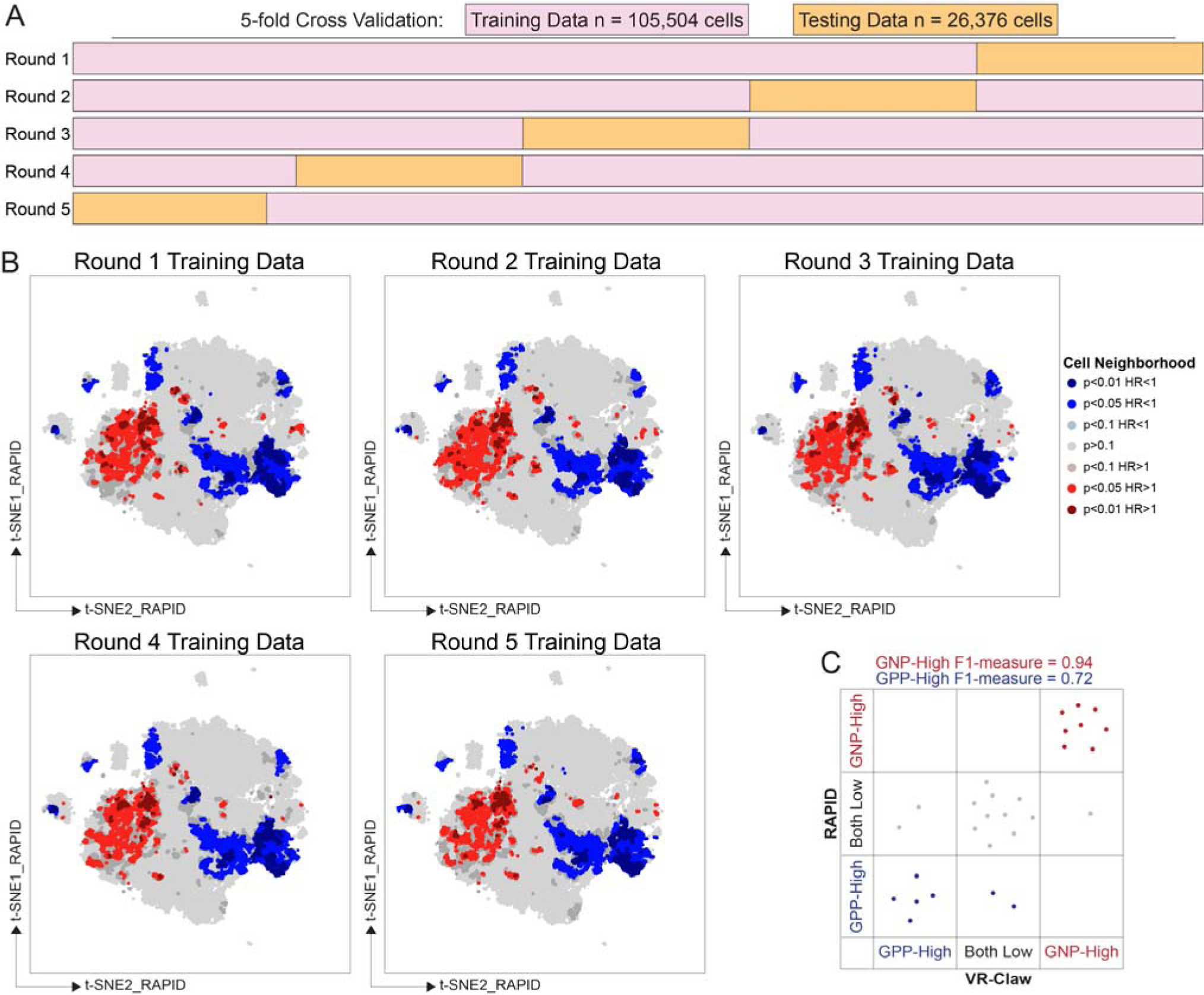
Velociraptor-Claw reproducibly identifies prognostic cells and classifies patients in Dataset 1. **A)** VR-Claw analyses of CD45^-^CD31^-^ GBM cells across each round of 5-fold cross validation. Cell neighborhoods are color coded based on HR and p-value. **B)** Patient classification according to prognostic population abundance by RAPID and VR-Claw.

**Supplementary Figure S2.**
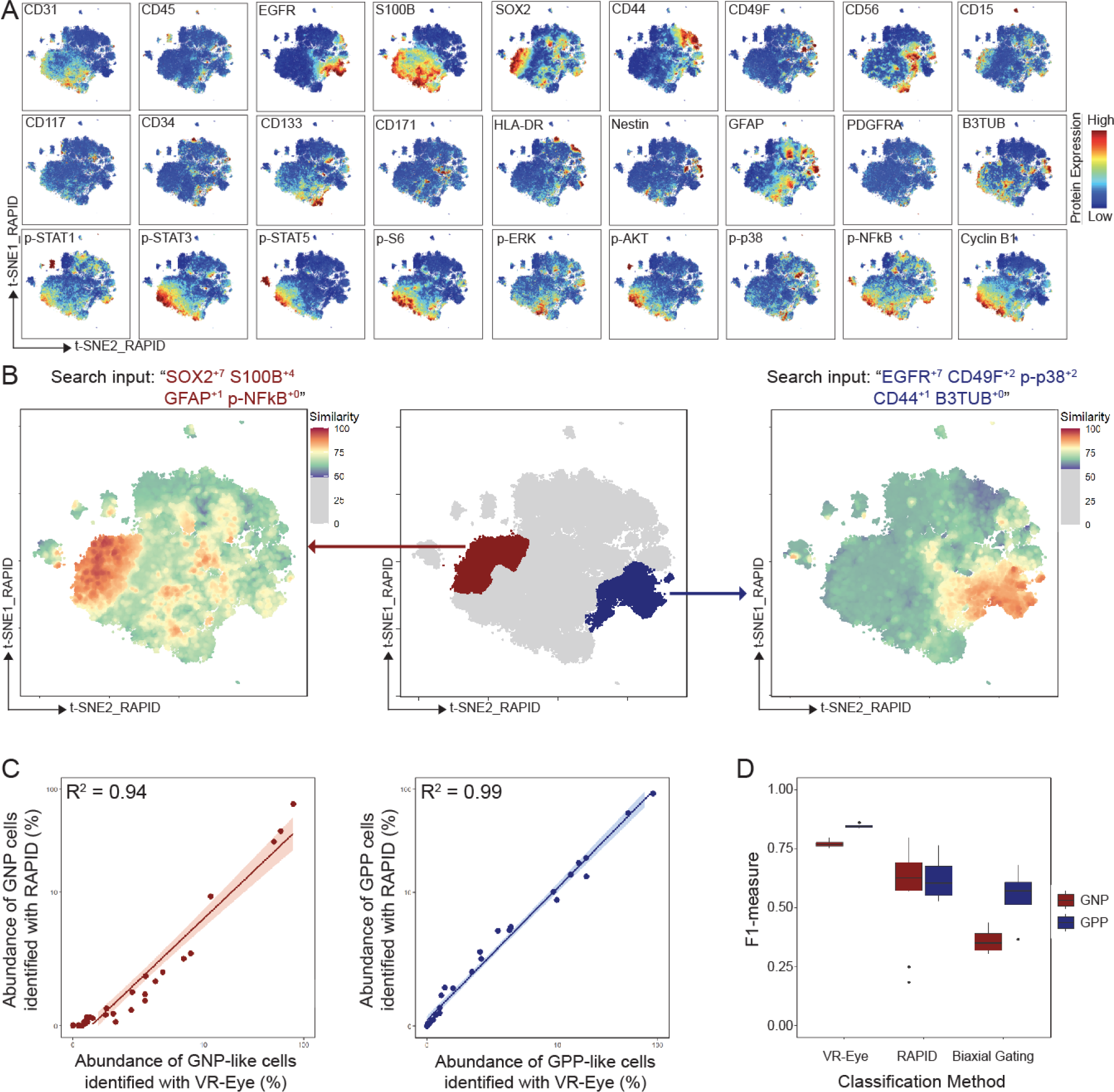
Optimization of prognostic GBM population identification. **A)** Expression levels of protein markers on common t-SNE axes for 131,880 CD45^-^CD31^-^ GBM cells in Dataset 1. A spectrum intensity scale indicates expression levels with blue representing low expression and red representing high expression. **B)** VR-Eye analysis seeking GNP-like cells (dark red in the middle plot) and GPP-like cells (dark blue in the middle plot). GNP-like cells, identified in the left plot, were sought with an input of SOX2^+7^ S100B^+4^ GFAP^+1^ p-NFkB^+0^). GPP-like cells, identified in the right plot, were sought with an input of EGFR^+7^ p-p38^+2^ CD49F^+2^ CD44^+1^ TUJ1^+0^) in the RAPID cohort (4,710 CD45^-^CD31^-^ GBM cells per patient). A spectrum intensity scale indicates cell similarity with red indicating high similarity and purple indicating low similarity. **C)** Correlation between each patient’s percentage of GNP-like or GPP-like cells, respectively, identified by RAPID and VR-Eye in Dataset 1. Pearson correlation tests were conducted. **D)** Comparison of cell identification approaches across multiple iterations of each approach. For VR-Eye and RAPID, each algorithm was applied to 10 different t-SNEs generated from the same sampling of Dataset 1 to identify GNP-like and GPP-like populations. Biaxial gating was performed by four different cytometry experts.

**Supplementary Figure S3.**
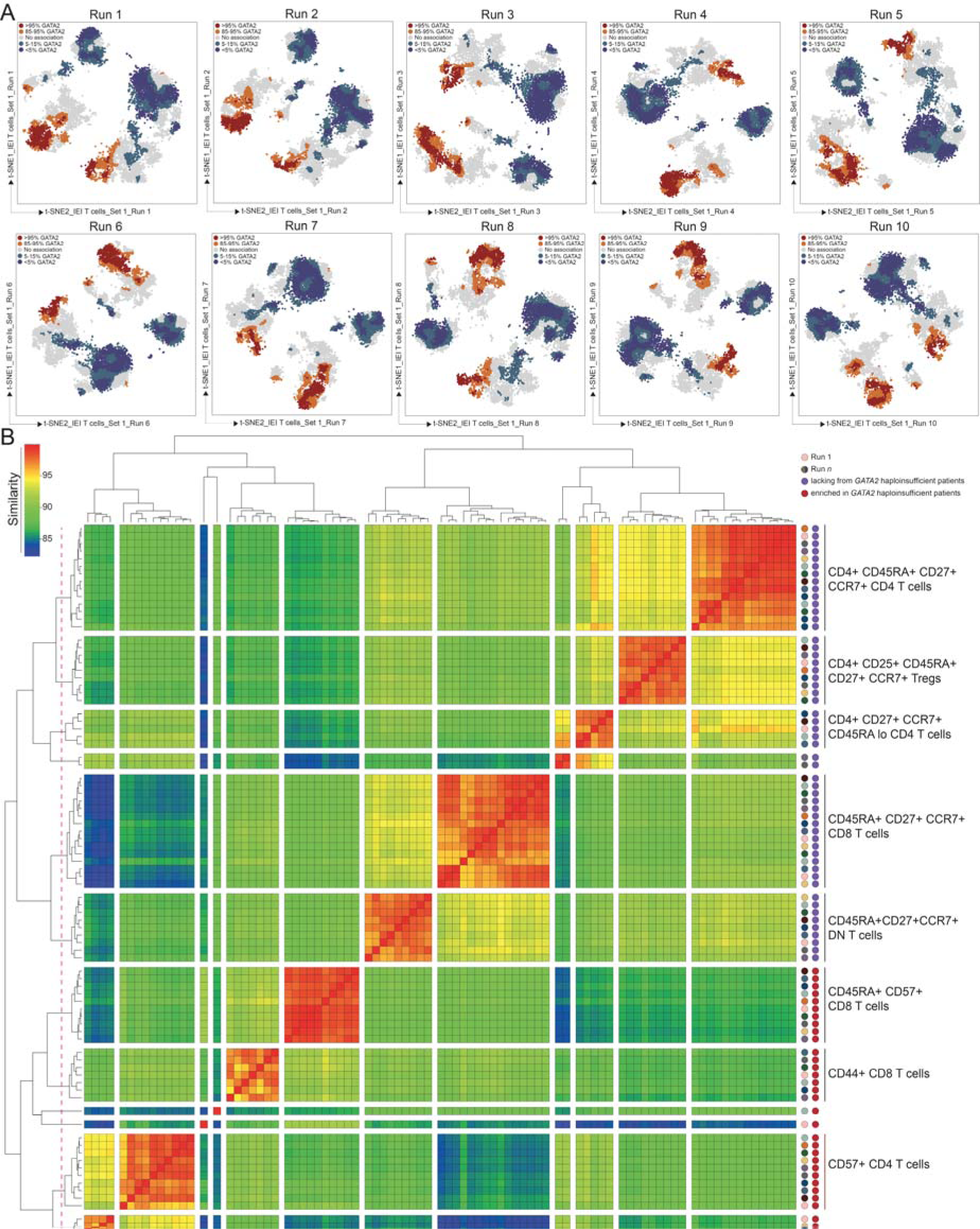
VR-Claw reproducibly reveals *GATA2* haploinsufficiency-specific cells in Dataset 3. **A)** VR-Claw analyses across 10 runs using 2,000 randomly sampled T cells per donor in Set 1 (N=5, n=10,000 cells total). Each t-SNE was independently generated for each run using that run’s sample of T cells. Red and orange denote populations enriched in *GATA2* haploinsufficiency patients; blue and purple denote populations lacking from *GATA2* haploinsufficiency patients. **B)** RMSD analysis on MEM labels generated from *GATA2* haploinsufficiency-specific populations identified in each of the 10 runs of VR-Claw. Similarity values were calculated by subtracting normalized RMSD values from 100. A rainbow intensity scale indicates population similarity with red indicating high similarity and purple indicating low similarity. Stable populations that appear in at least 5 out of 10 runs are marked to the right of the heatmap with either a blue or a dark red line. Blue lines indicate populations that were lacking from *GATA2* haploinsufficiency patients, and dark red lines indicate populations that were enriched in *GATA2* haploinsufficiency patients as determined by VR-Claw. Phenotypes of stable populations are summarized to the far right.

**Supplementary Figure S4.**
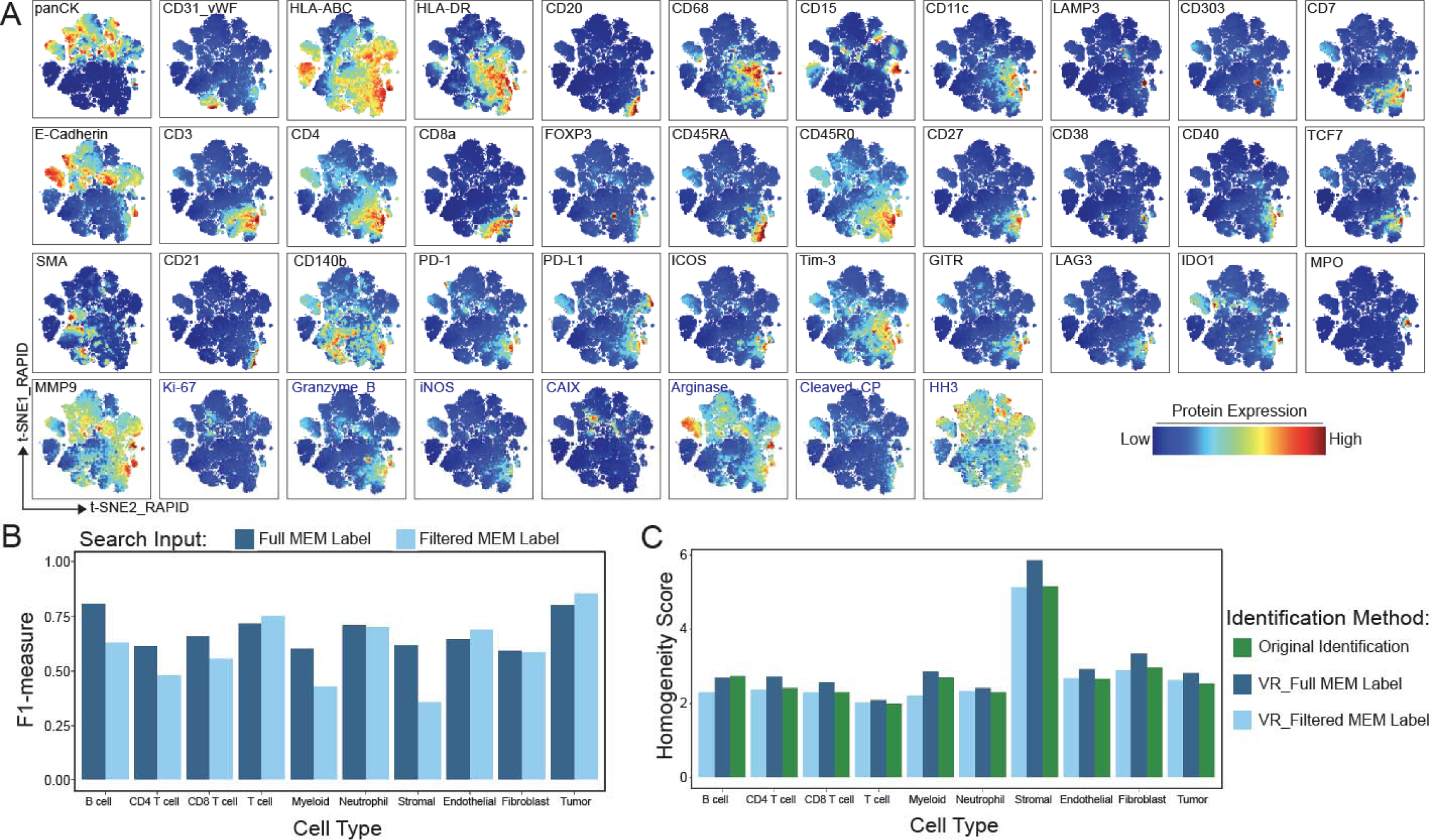
41-dimensional phenotypic labels outperform filtered phenotypes in Dataset 4. **A)** Expression levels of protein markers on common t-SNE axes for 257,076 cells in Dataset 4. A spectrum intensity scale indicates expression levels with blue representing low expression and red representing high expression. Markers not included as t-SNE input are labeled in blue. **B)** Comparison of VR-Eye cell identification using full 41-dimensional MEM labels or filtered MEM labels (filtered labels range from 4- dimensional to 28-dimensional) based on population protein enrichment as search input. **C)** Comparison of population homogeneity across different cell identification methods. The homogeneity score equals the inverse of the median RMSD with respect to the median protein expression for each marker for each population.

**Supplementary Figure S5.**
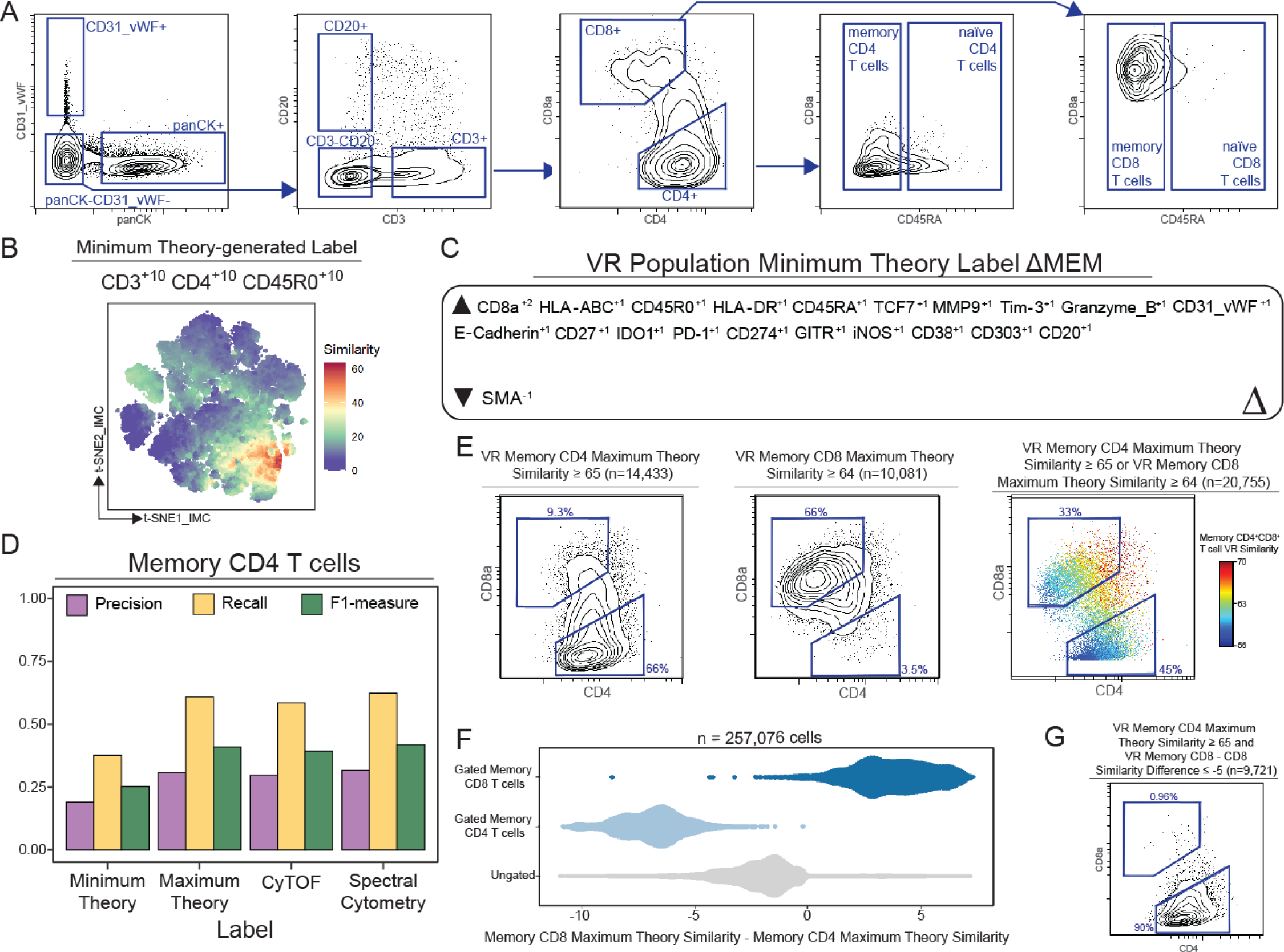
Velociraptor identifies CD4^+^CD8^+^ double positive memory T cells present in human breast cancer tumors. **A)** Gating scheme to select memory CD4 T cells shown on a representative patient. **B)** VR-Eye analysis seeking memory CD4 T cells using a minimum marker label. **C)** ΔMEM analysis comparing the VR-identified memory CD4 T cell population to the gated population. ΔMEM labels were calculated by subtracting the absolute MEM label of the VR-identified population from the absolute MEM label of the manually gated population. **D)** Quantification of search accuracy for each memory CD4 T cell label sought. Precision is plotted in purple, recall is plotted in gold, and F1-measures are plotted in green. Similarity thresholds were optimized to best capture the gated population. **E)** Biaxial plots of CD4 and CD8a expression are shown for cells included in Velociraptor-identified memory CD4 (left plot) or CD8 (middle plot) populations. Contour indicates density. A spectrum intensity scale indicates each cell’s similarity to the memory CD4^+^CD8^+^ double positive reference phenotype (right plot) as determined by maximum theory searches. **F)** Density plots show the difference between each cell’s memory CD4 similarity and memory CD8 similarity to the maximum theory label. Biaxially gated memory CD4 T cells are shown in light blue, and biaxially gated memory CD8 T cells are shown in dark blue. Ungated cells are shown in light grey. **G)** Biaxial plots of CD4 and CD8a expression are shown for cells with a memory CD4 T cell similarity value greater than or equal to 65 and a memory CD8 – CD4 similarity difference less than or equal to -5. Contour indicates density.

