## Supplementary Tables for "Velociraptor: Cross-Platform Quantitative Search Using Hallmark Cell Features"

Supplementary Table 1: Recurrent GBM Mass Cytometry Panel for Dataset 2

| Metal | Mass | Marker | Clone | Vendor | Catalog Number | Condition | Dilution |
|---|---|---|---|---|---|---|---|
| Rh | 103 | Live/Dead | - | Fluidigm | 201103A | extracellular | - |
| La | 139 | Cyclin B1 | GNS-1 | BD Biosciences | 554176 | methanol | 1:100 |
| Pr | 141 | TUBB3 | TUJ1 | Biolegend | 801201 | methanol | 1:100 |
| Nd | 142 | cCasp3 | 5AE1 | Fluidigm | 3142004A | methanol | 1:100 |
| Nd | 143 | CD117 | 104D2 | Fluidigm | 3143001B | extracellular | 1:100 |
| Nd | 144 | S100B | 19/S100B | BD Biosciences | 612376 | methanol | 1:100 |
| Nd | 145 | CD31 | WM59 | Fluidigm | 3145004B | extracellular | 1:100 |
| Sm | 147 | gH2AX | JBW301 | Fluidigm | 3147016A | methanol | 1:100 |
| Nd | 148 | CD34 | 581 | Fluidigm | 3148001B | extracellular | 1:100 |
| Sm | 149 | p-4EBP1 (T37/T46) | 236B4 | Fluidigm | 3149005A | methanol | 1:100 |
| Nd | 150 | p-STAT5 (Y694) | 47 | Fluidigm | 3150005A | methanol | 1:100 |
| Eu | 151 | BMX | 40/BMX | BD Biosciences | 610793 | methanol | 1:100 |
| Sm | 152 | p-AKT (S473) | D9E | Fluidigm | 3152005A | methanol | 1:100 |
| Eu | 153 | p-STAT1 (Y701) | 58D6 | Fluidigm | 3153003A | methanol | 1:100 |
| Sm | 154 | CD45 | HI30 | Fluidigm | 3154001B | extracellular | 1:400 |
| Gd | 155 | CD56 (NCAM) | HCD56 | Biolegend | 318302 | extracellular | 1:100 |
| Gd | 156 | p-p38 (T180/Y182) | D3F9 | Fluidigm | 3156002A | methanol | 1:100 |
| Gd | 158 | p-STAT3 (Y705) | 4/P-STAT3 | Fluidigm | 3158005A | methanol | 1:100 |
| Tb | 159 | CD49F | GoH3 | Biolegend | 313602 | extracellular | 1:100 |
| Dy | 161 | PDGFRa | 16A1 | Biolegend | 323502 | extracellular | 1:50 |
| Dy | 163 | SOX2 | O30-678 | BD Biosciences | 561469 | saponin | 1:100 |
| Dy | 164 | CD15 | W6D3 | Fluidigm | 3164001B | extracellular | 1:100 |
| Ho | 165 | EGFR | AY13 | Biolegend | 352902 | extracellular | 1:100 |
| Er | 166 | p-NFkB (S529) | K10-895.12.50 | Fluidigm | 3166006A | methanol | 1:100 |
| Er | 167 | L1CAM | 5G3 | BD Biosciences | 554273 | extracellular | 1:100 |
| Er | 168 | Nestin | 10C2 | Millipore | MAB5326 | methanol | 1:100 |
| Tm | 169 | CD44 | BJ18 | Biolegend | 338802 | extracellular | 1:100 |
| Er | 170 | GFAP | 1B4 | BD Biosciences | 556328 | methanol | 1:200 |
| Yb | 171 | p-ERK1/2 (T202/Y204) | D13.14.4E | Fluidigm | 3171010A | methanol | 1:100 |
| Yb | 172 | p-S6 (S235/S236) | N7-548 | Fluidigm | 3172008A | methanol | 1:100 |
| Yb | 173 | SOX10 | A-2 | Santa Cruz | sc-365692 | methanol | 1:100 |
| Yb | 174 | HLA-DR | L243 | Fluidigm | 3174001B | extracellular | 1:200 |
| Lu | 175 | p-HH3 | HTA28 | Fluidigm | 3175012A | methanol | 1:400 |
| Yb | 176 | HH3 | D1H2 | Fluidigm | 3176016A | methanol | 1:200 |
| Ir | 191 | Intercalator | - | Fluidigm | 201192B | methanol | - |
| Ir | 193 | | | | | methanol | |

Supplementary Table 2: T Cell Immunophenotyping Mass Cytometry Panel for Dataset 3

| Metal | Mass | Marker | Clone | Vendor | Catalog Number | Condition | Dilution | t-SNE to select T cells | Donor-specific t-SNE |
|---|---|---|---|---|---|---|---|---|---|
| Y | 89 | CD45 | HI30 | Fluidigm | 3089003B | extracellular | 1:200 | X | |
| Rh | 103 | Live/Dead | - | Fluidigm | 201103B | extracellular | - | | |
| Pd/Cd | 106 | CD66b | 80H3 | Biolegend | 305102 | extracellular | 1:100 | X | |
| Pd/Cd | 110 | CD16 | 3G8 | Biolegend | 302051 | extracellular | 1:100 | X | |
| Cd | 111 | CD8 | RPAT8 | Biolegend | 301053 | extracellular | 1:100 | X | X |
| Cd | 112 | CD14 | M5E2 | Biolegend | 301843 | extracellular | 1:100 | X | |
| Cd | 113 | CD4 | RPA-T4 | Biolegend | 300541 | extracellular | 1:100 | X | X |
| Cd | 114 | CD3 | UCHT1 | Biolegend | 300443 | extracellular | 1:100 | X | |
| Cd | 116 | CD19 | HIB19 | Biolegend | 302247 | extracellular | 1:100 | X | |
| Pr | 141 | CD45R0 | UCHL1 | Biolegend | 304239 | extracellular | 1:100 | X | X |
| Nd | 142 | CPT1a | 8F6AE9 | Abcam | ab128568 | methanol | 1:100 | | |
| Nd | 143 | CD127 | A019D5 | Fluidigm | 3143012B | extracellular | 1:200 | X | X |
| Nd | 144 | ATP5a | 7H10BD4F9 | Abcam | ab110273 | methanol | 1:100 | | |
| Nd | 145 | GRIM19 | 6E1BH7 | Abcam | ab110240 | methanol | 1:100 | | |
| Sm | 147 | CD20 | 2H7 | Fluidigm | 3147001B | extracellular | 1:200 | X | |
| Nd | 148 | CD27 | L128 | BD | 624084 | extracellular | 1:100 | X | X |
| Sm | 149 | CCR4 | 205410 | Fluidigm | 3149029A | extracellular | 1:200 | X | X |
| Nd | 150 | CD134 | ACT35 | Fluidigm | 3150023C | extracellular | 1:100 | X | X |
| Eu | 151 | ICOS | C398.4A | Fluidigm | 3151020B | extracellular | 1:200 | X | X |
| Sm | 152 | TCRgd | 11F2 | Fluidigm | 3152008B | extracellular | 1:200 | X | X |
| Sm | 154 | GLUT3 | polyclonal | Abcam | ab41525 | methanol | 1:100 | | |
| Gd | 156 | CXCR3 | G025H7 | Fluidigm | 3156004B | extracellular | 1:200 | X | X |
| Gd | 158 | CD137 | 4B4-1 | Fluidigm | 3158013B | extracellular | 1:200 | X | X |
| Tb | 159 | CCR7 | G043H7 | Fluidigm | 3159003A | extracellular | 1:100 | X | X |
| Gd | 160 | CD98 | MEM-108 | Biolegend | 315602 | extracellular | 1:100 | | |
| Dy | 161 | CTLA4 | 14D3 | Fluidigm | 3161004B | methanol | 1:100 | X | X |
| Dy | 162 | Ki-67 | B56 | Fluidigm | 3162012B | methanol | 1:200 | | |
| Dy | 163 | GLUT1 | polyclonal | Novus Biologicals | NB110-39113 | methanol | 1:100 | | |
| Dy | 164 | CD95 | DX2 | Fluidigm | 3164008B | extracellular | 1:200 | X | X |
| Er | 166 | CD44 | BJ18 | Fluidigm | 3166001C | extracellular | 1:100 | X | X |
| Er | 167 | CD38 | HIT2 | Fluidigm | 3167001B | extracellular | 1:100 | X | X |
| Er | 168 | CYTOC | 6H2.B4 | BD | 556432 | methanol | 1:100 | | |
| Tm | 169 | CD25 | 2A3 | Fluidigm | 3169003B | extracellular | 1:200 | X | X |
| Er | 170 | CD45RA | HI100 | Fluidigm | 3170010B | extracellular | 1:200 | X | X |
| Yb | 171 | CXCR5 | RF8B2 | Fluidigm | 3171014B | extracellular | 1:200 | X | X |
| Yb | 172 | CD57 | HCD57 | Fluidigm | 3172009B | extracellular | 1:100 | X | X |
| Yb | 173 | CXCR4 | 12G5 | Fluidigm | 3173001B | extracellular | 1:200 | X | X |
| Yb | 174 | HLA-DR | II L243 | Fluidigm | 3174001B | extracellular | 1:200 | X | X |
| Lu | 175 | PD-1 | EH12.2H7 | Fluidigm | 3175008B | extracellular | 1:200 | X | X |
| Yb | 176 | CD56 | CMSSB | Fluidigm | 3176003B | extracellular | 1:200 | X | X |
| Ir | 191 | Intercalator | - | Fluidigm | 201192B | methanol | - | | |
| Ir | 193 | | | | | | | | |
| Bi | 209 | CD11b | ICRF44 | Fluidigm | 3209003B | extracellular | 1:200 | X | |
